# Particle flow modulates growth dynamics and nanoscale-arrested growth of transcription factor condensates in living cells

**DOI:** 10.1101/2022.01.11.475940

**Authors:** Gorka Muñoz-Gil, Catalina Romero-Aristizabal, Nicolas Mateos, Felix Campelo, Lara I. de Llobet Cucalon, Miguel Beato, Maciej Lewenstein, Maria F. Garcia-Parajo, Juan A. Torreno-Pina

**Affiliations:** ICFO-Institut de Ciencies Fotoniques, The Barcelona Institute for Science and Technology (BIST), 08860 Barcelona, Spain; Centre de Regulació Genomica (CRG), The Barcelona Institute of Science and Technology (BIST), Dr. Aiguader 88, Barcelona, Spain; Universitat Pompeu Fabra (UPF), Barcelona, Spain; ICREA, Pg. Lluis Companys 23, 08010 Barcelona, Spain

**Keywords:** Biomolecular condensates, Brownian motion coalescence, Liquid-liquid phase separation, Single particle tracking, Machine learning.

## Abstract

Liquid-liquid phase separation (LLPS) is emerging as key physical principle for biological organization inside living cells, forming condensates that play important roles in the regulation of multiple functions. Inside living nuclei, transcription factor (TF) condensates regulate transcriptional initiation and amplify transcriptional output of expressed genes. Yet, the biophysical parameters controlling TF condensation are still poorly understood. Here we applied a battery of single molecule imaging tools, theory and simulations to investigate the physical properties of TF condensates of the Progesterone Receptor (PR) *in vivo*. Analysis of individual PR trajectories at different ligand concentrations showed marked signatures of a ligand-tunable and regulated LLPS process. Using a machine learning architecture, we uncovered that diffusion within condensates follows fractional Brownian motion, reflecting viscoelastic interactions between PR and chromatin within condensates. High density single molecule localization maps further revealed that condensate growth dynamics is dominated by Brownian motion coalescence (BMC) at shorter times, but deviate at longer timescales reaching a growth plateau with nanoscale condensate sizes. To understand our observations we developed an extension of the BMC model by including stochastic unbinding of particles within condensates. The model reproduced the BMC behavior together with finite condensate sizes a steady-state, fully recapitulating our experimental data. Our results are thus consistent with droplet growth dynamics being regulated by the escaping probability of TFs molecules from condensates. The interplay between condensation assembly and molecular escaping maintains an optimum physical condensate size. Such phenomena must have implications for the biophysical regulation of other TF condensates and could also operate in multiple biological scenarios.

## Introduction

Activities performed by living cells are generally achieved through the compartmentalization of their multiple components in space and time. Although traditionally cell compartments have been thought to be surrounded by membranes, recent evidence indicate that cells also organize membrane-less internal compartments through the physical process of liquid-liquid phase separation (LLPS)(1-4). LLPS creates transient chemically distinct compartments, also called biomolecular condensates, which might operate as versatile biochemical “hubs” inside the cell(1, 5). Phase separation is particularly relevant in the cell nucleus, where the condensation of numerous proteins on chromatin have been shown to regulate gene transcription and chromatin architecture at multiple temporal and spatial scales(6-8). Transcription factor (TF) condensates are proposed to regulate transcriptional initiation and amplify transcriptional output of expressed genes(5, 7, 9-11). Yet, despite its prevalence and biological significance, quantitative determination and understanding of the biophysical parameters controlling TF condensation in the nucleus of living cells is largely missing.

Nuclear receptors are a family of TFs that have been widely studied as master regulators of gene transcription and genome topology in response to an external stimulus: a steroid hormone(12-14). Structurally, these TFs contain two intrinsically disordered regions that favor phase separation: the N-terminal domain and the hinge; as well as two highly structured regions: the DNA-binding domain and the ligand-binding domain(15). Ligand stimulation of several members of this family has been shown to trigger LLPS, forming nuclear condensates with different transcriptional roles(16-18). Since ligand addition allows accurate control of the onset for nucleation and condensate coarsening, nuclear receptors represent an ideal system to study inducible phase separation and to follow their temporal evolution in well-controlled and tunable experimental settings.

From the theoretical side, phase separation is usually associated to the heterogeneous mixing of two components, either by spinodal decomposition (19), or nucleation(20). In general, entropy based models, as e.g. the Flory-Huggins model (21, 22), have been commonly used to understand phase separated systems in biological scenarios(23). Moreover, in recent years, several studies have addressed the temporal evolution of condensate nucleation and growth within the full complexity of living cells. For instance, it has been shown that biocondensate nucleation and coarsening can be described by different physical mechanisms such as diffusion-limited growth, diffusion Ostwald ripening (OR), or Brownian motion coalescence (BMC)(24). The common physical property underlying these mechanisms is a dynamic power-law scaling behavior of the mean droplet sizes(24), with a final steady-state that results in a single condensate containing all phase separated molecules. However, consistent deviations from these LLPS growing mechanisms have been also reported and attributed to the occurrence of active nonequilibrium processes within living cells, such as RNA transcription(24) or the presence of obstacles such as chromatin(25). Hence, models able to predict and/or adapt classical phase separation properties to the living cell context are still under development.

Here we investigate the physical properties of LLPS in transcriptional condensates of the nuclear Progesterone Receptor (PR) *in vivo* using an extensive combination of single molecule approaches, theory and simulations. Analysis of single PR trajectories showed a hormone-dependent bimodal distribution on the diffusion of the receptor associated to particles diffusing within and outside condensates. Using a deep learning method, we uncovered that diffusion within condensates is best described by means of fractional Brownian motion(26). High density single molecule localization maps as function of time further revealed a BMC-like growth process at shorter times, but that markedly deviated at longer timescales reaching a growth plateau on the condensate sizes at the nanoscale. To quantitatively understand our observations we developed an extension of the BMC model by including stochastic unbinding of particles within condensates. Our model is not only able to reproduce the usual BMC behavior, but importantly, it also reaches a steady-state with finite condensate sizes. As a whole, our single molecule experimental data and theoretical model is consistent with droplet growth dynamics being regulated by the escaping probability of TFs molecules from condensates.

### Single Particle Tracking of nuclear PR *in vivo* in response to a tunable stimulus

As most nuclear receptors, PR contains an intrinsically disorder N-terminal domain region (Fig. S1) and thus, it is prone to phase separate. We first confirmed LLPS of PR in the nucleus of living breast cancer cells after hormone exposure using confocal microscopy. Condensates visibly formed minutes after adding the hormone (Video S1). However and contrary to a vast literature in the field, PR condensates remained relatively small in size, being clearly diffraction-limited. We thus turned to single molecule approaches to effectively increase the spatial (∼20 nm) and temporal (∼15 ms) resolution providing dynamic information on the behavior of individual PR molecules in the nucleus. In particular, we applied single particle tracking (SPT) which has been widely used over the last decade to evaluate the lateral mobility of several TFs and DNA binding proteins in the nucleus of living cells at the single molecule level(13, 27-30). We generated a stable MCF7 breast cancer cell line expressing a SNAP-GFP-PRB (PR isoform B)(31). PR molecules were labeled with the SNAP-JaneliaFluor 549 (JF549) dye (32) and their diffusion inside the nucleus of living cells was recorded under highly inclined illumination at a frame rate of 15ms, as schematically illustrated in Fig. 1A. Individual JF549 localizations were reconnected to generate trajectories that were analyzed by computing the time averaged mean square displacement (tMSD) and the angular distribution over consecutive steps as shown in Fig. 1B (28, 29). The instantaneous diffusion coefficients for each trajectory were extracted by linear fitting of the 2^nd^ and 4^th^ points (*D*_2−4_) of the tMSD curve(33) and used to build up D_2−4_histograms of hundreds of trajectories over different cells. In addition, the angular distribution provides information on the type of diffusion exhibited by a molecule while interacting with its environment. Whereas the angular distribution is uniform when molecules diffuse in a homogeneous environment, an asymmetric angular distribution with a preferred occurrence of angles at 180° reflects the presence of confinement or obstacles to the molecule diffusion(29).

**Figure 1.**
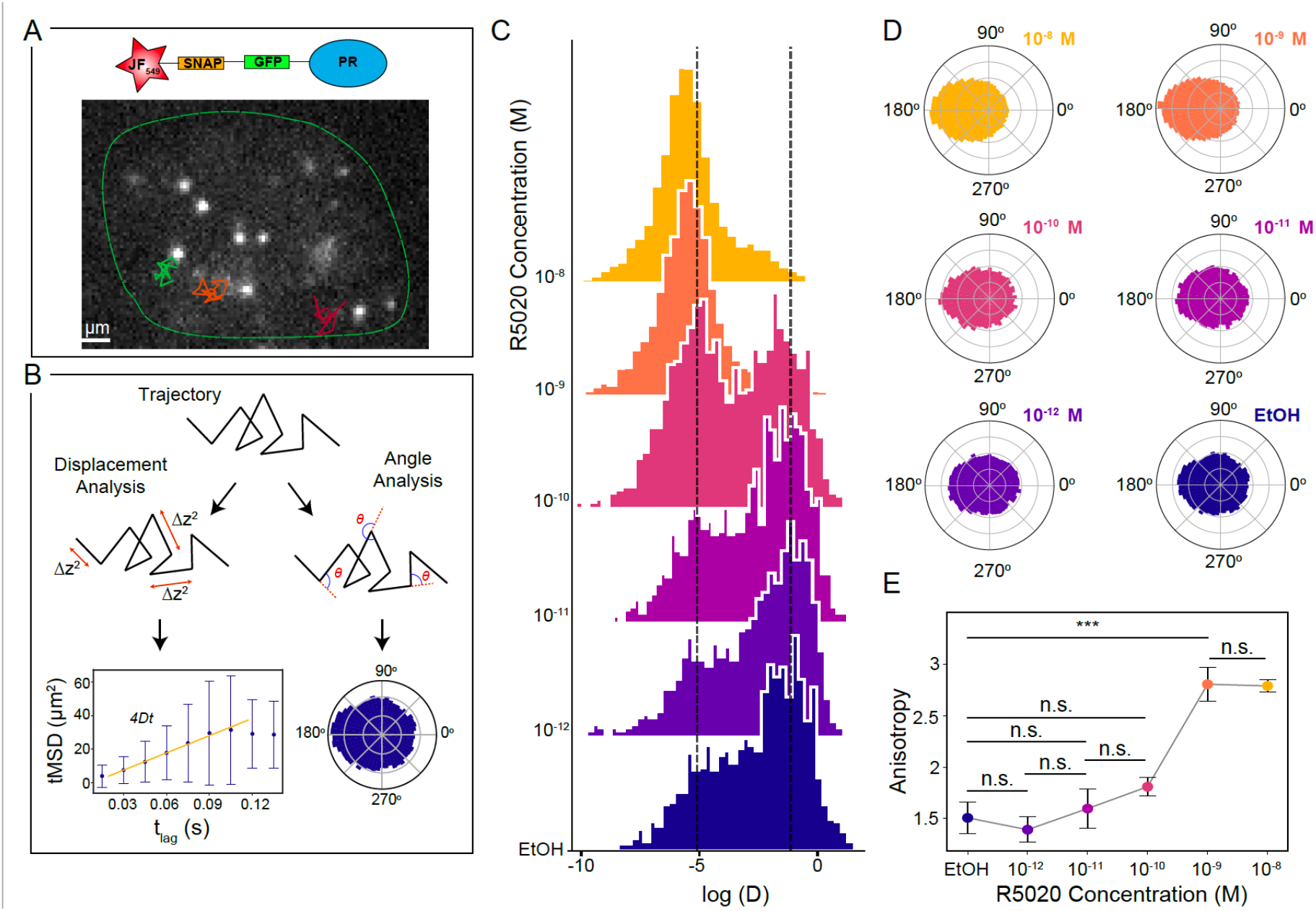
Lateral diffusion of individual PR molecules in the nucleus of living cells. (A) Representative frame of a SPT video. Individual PR molecules (bright spots) were visualized in the nucleus (green outline) of MCF7 breast cancer cells, under a highly inclined illumination at 15 ms frame rate. Diffraction-limited single molecule localizations were tracked in successive frames to generate individual trajectories (super-imposed color lines). (B) Schematic representation of the trajectory analysis. For each trajectory we extract the displacement between frames to generate individual tMSD plots as a function of the time lag and extract the diffusion coefficients (*D*_2−4_) for each trajectory (left, lower panel). In addition, we calculate the angles between successive steps to create polar histograms (right, lower panel). (C) Distribution of the *D*_2−4_(μm^2^/s) values of individual PR trajectories exposed to increasing R5020 concentrations for 1 hour. EtOH corresponds to the control condition, i.e., in the absence of ligand. Y axis corresponds to the frequency of events. Vertical dash lines indicate *D*_2−4_values 0.0061 (left line) and 0.5 μm^2^/s (right line). Data extracted from at least 1000 trajectories belonging to at least 8 cells from 3 independent experiments. (D) Polar histograms of the angle between successive steps of diffusing PR under increasing R5020 concentrations. (E) Anisotropy values as a function of R5020 concentration for at least 8 cells analyzed. Results of a one-way analysis of variance (ANOVA) test are shown as: n.s. for not significant, *** for p-value< 0.001.

To investigate the PR lateral mobility in response to hormone, MCF7 cells were treated with a broad range of concentrations of the progesterone derivative R5020 (10^−12^ M to 10^−8^ M, for 1 hour), or with EtOH as a control(34). As shown in Fig. 1C, we mainly observe two populations in the distribution of *D*_2−4_values across different concentrations, similar to other proteins that interact with chromatin(27, 28). Strikingly, instead of a gradual increase in the bound fraction of PRs that one would expect from a stochiometric occupancy of TFs to DNA binding sites with increasing ligand concentration, we found a sharp transition from freely to bound fraction taking place at a critical ligand concentration of 10^−10^ M (Fig. 1C). This sharp transition in PR mobility suggests that LLPS might be regulating the interaction between PR and chromatin.

We further computed the distribution of angles between consecutive displacements for each individual trajectory on multiple cells and for different hormone concentrations. At hormone concentrations 10^−10^ M and below, diffusion is mainly isotropic and PR explores all angles with equal probability (Fig. 1D). In strong contrast, above the critical concentration of 10^−10^ M, the angle distributions become highly anisotropic with an increased occurrence of angles at 180°, i.e., higher probability for PR molecules to bounce back to their prior position (Fig. 1D). To better quantify these results, we computed the degree of anisotropy as the fold increase of angles occurring at 180° ± 30° with respect to 0°± 30° (28). A sharp transition in anisotropy was retrieved above 10^−10^ M R5020 concentration (Fig. 1E), alike to that at which the *D*_2−4_sharp transition takes place. We interpret this preferential backward movement as evidence of confinement, and an indication of the bias in angles experienced by a particle inside a condensate when being constrained by the condensate boundaries. Altogether, our SPT results are consistent with a ligand-tunable and regulated LLPS process.

### Diffusion behavior of individual PR determined with machine learning

Due to the short length of the SPT trajectories (usually less than 30 time segments), it is challenging to identify the diffusion behavior of PR inside living nuclei using conventional data analysis methods. We thus relied on a recently developed machine learning (ML) analysis(35). Using a combination of convolutional and recurrent neural networks (see Methods, Fig. S2) we: (1) identified the theoretical model that best describes the diffusion behavior of individual PR trajectories, and (2) determined the corresponding anomalous exponent *α*, defined as the scaling factor when fitting the tMSD to a power-law ∼ *tα*. Here, *α* = 1 corresponds to Brownian diffusion, *α* < 1 to anomalous sub-diffusion, and *α* > 1 to super-diffusion. We first trained the algorithm with a set of simulated trajectories arising from various diffusion models related to many different experimental observations (see Methods). Remarkably, when applied to our single molecule experimental data, the ML algorithm revealed two main types of diffusion (Fig. S3) i.e., the vast majority of the trajectories were either classified as diffusing according to the annealed transit time model (ATTM)(36), or exhibiting fractional Brownian motion (FBM)(37). ATTM has been associated to the anomalous, non-ergodic and non-Gaussian motion of particles diffusing in a spatiotemporal heterogeneous medium, e.g. on cell membranes(38). FBM has been described as an extension of Brownian motion with correlated noise, often associated to diffusion in viscoelastic media(39). Note that, since the trajectories were normalized before entering the ML architecture (see Methods), the ML prediction is independent of the diffusion coefficient value. For each hormone concentration, we computed the percentage of trajectories predicted as ATTM or FBM. At ligand concentrations below 10^−10^ M, 60% of the trajectories were classified as ATTM and 40% as exhibiting FBM (Fig. 2A). Notably, a sharp change in the diffusion behavior occurs at > 10^−10^ M R5020, with ∼ 80% of the trajectories exhibiting FBM and ∼ 20% ATTM (Fig. 2A). We further exploited the powerful discrimination capability of the ML algorithm to compute the *D*_2−4_values of the trajectories assigned to each of the theoretical models. We found that FBM trajectories display a much lower lateral mobility as compared to those assigned to ATTM (Fig. 2B). Together, the sharp increase in the number of molecules exhibiting FBM together their lower mobility at ligand concentrations above 10^−10^ M suggest that PR diffusion behavior results from viscoelastic interactions between the receptor and chromatin within a condensate.

**Figure 2.**
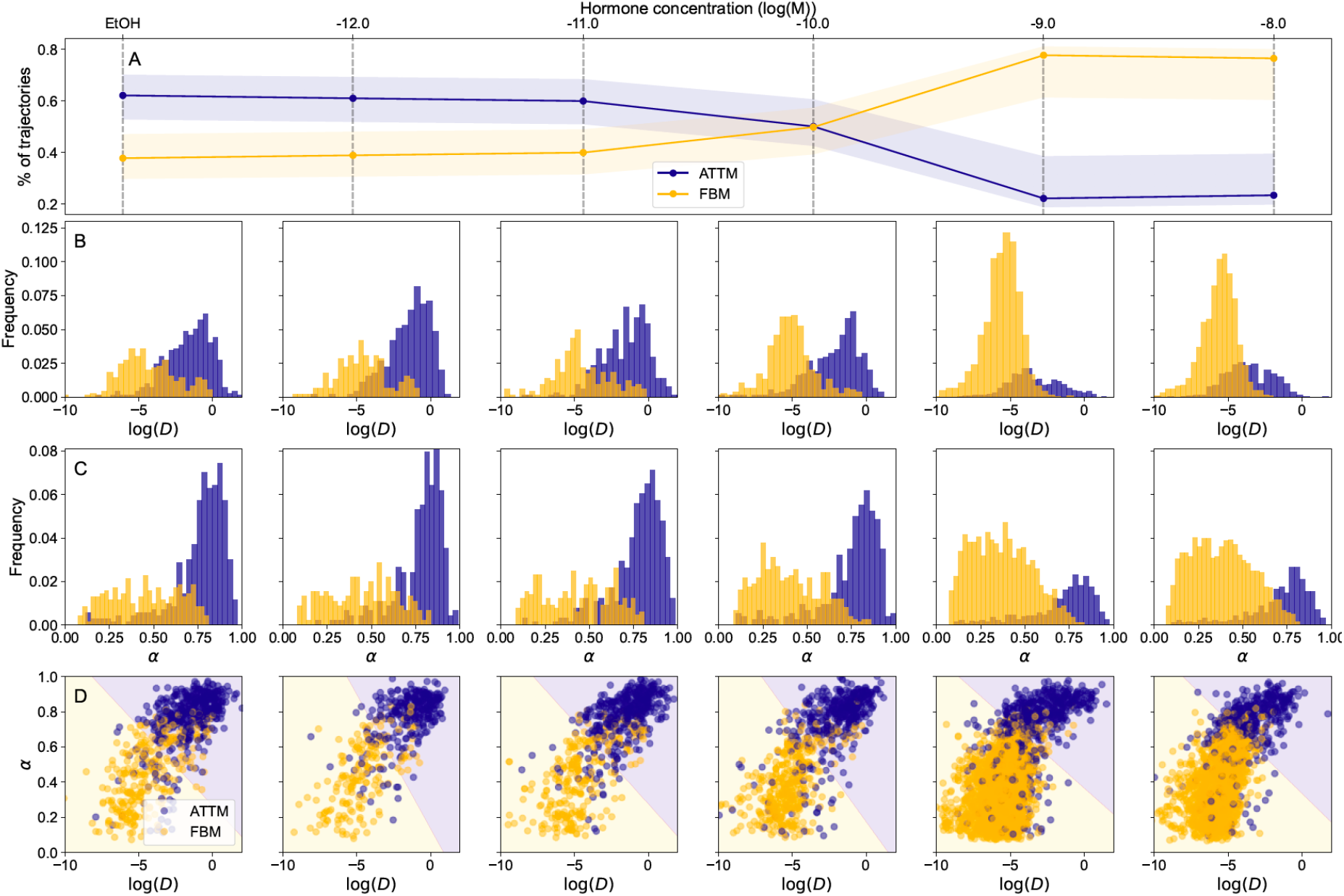
Machine learning analysis of individual PR trajectories in living cells. (A) Percentage of trajectories associated to ATTM (blue) or FBM (yellow) by the ML algorithm as a function of ligand concentration. The shadow areas represent the error of the prediction, calculated by means of a confusion matrix (see Methods) (B) *D*_2−4_(μm^2^/s) distributions for varying ligand concentrations, with trajectories associated to ATTM (blue) and FBM (yellow), as identified by ML. (C) Corresponding histograms of the ML predicted anomalous exponents. (D) Scatter plot of the *D*_2−4_vs. anomalous exponent for every trajectory. Background color represents the prediction of a SVM trained on the data (see Methods).

Using a different ML architecture as described in Methods, we also predicted the *α* values for each of the observed trajectories. We found that FBM trajectories exhibit on average lower *α* values (∼ 0.45) than ATTM trajectories (∼ 0.75) (Fig. 2C). To assess the relationship between *D* and *α*, we generated scatterplots of these two parameters for different ligand concentrations (Fig. 2D). Strikingly, trajectories assigned to either ATTM or FBM form two differentiated clusters that can be readily classified by a support vector machine (SVM). The background color used in Fig. 2D shows the predictions of the SVM, demonstrating that *D* and *α* are sufficient to separate the lateral diffusion behavior of individual PRs as a function of ligand concentration. Overall, the ML analysis accurately separates two PR populations diffusing in markedly different media; and most importantly, it reflects a critical ligand concentration at which a transition from unbound (ATTM) to chromatin-bound (FBM) takes place.

### Nanometer-scale temporal evolution of PR condensates in living nuclei

Our single molecule mobility analysis is consistent with the emergence of PR condensates in living nuclei above a critical ligand concentration, but it does not provide direct information on the condensate sizes. To enquire on the relevant spatiotemporal scales involved in PR condensation and its temporal evolution we took advantage of the nanometer localization precision encoded in the SPT data. We used this information to generate 2D density maps of individual PR localization positions as the receptor dynamically explores the nuclear region(33). The 2D maps clearly show that hormone treatment (10^−8^ M, 60 min) leads to a strong accumulation of single molecule localization events in small regions, as compared to control conditions (Fig. 3A). We further evaluated the lateral mobility inside condensates by reconnecting the localization positions over consecutive frames. Remarkably, PR trajectories within condensates reproduce the mobility, angle distribution, and FBM diffusion behavior of the slow population retrieved by standard SPT shown in Figs. 1 and 2 (Fig. S4). These results also confirm that the slow population retrieved from the SPT analysis corresponds to the diffusion of PR molecules *inside* condensates rather than to the diffusion of the condensate itself.

**Figure 3.**
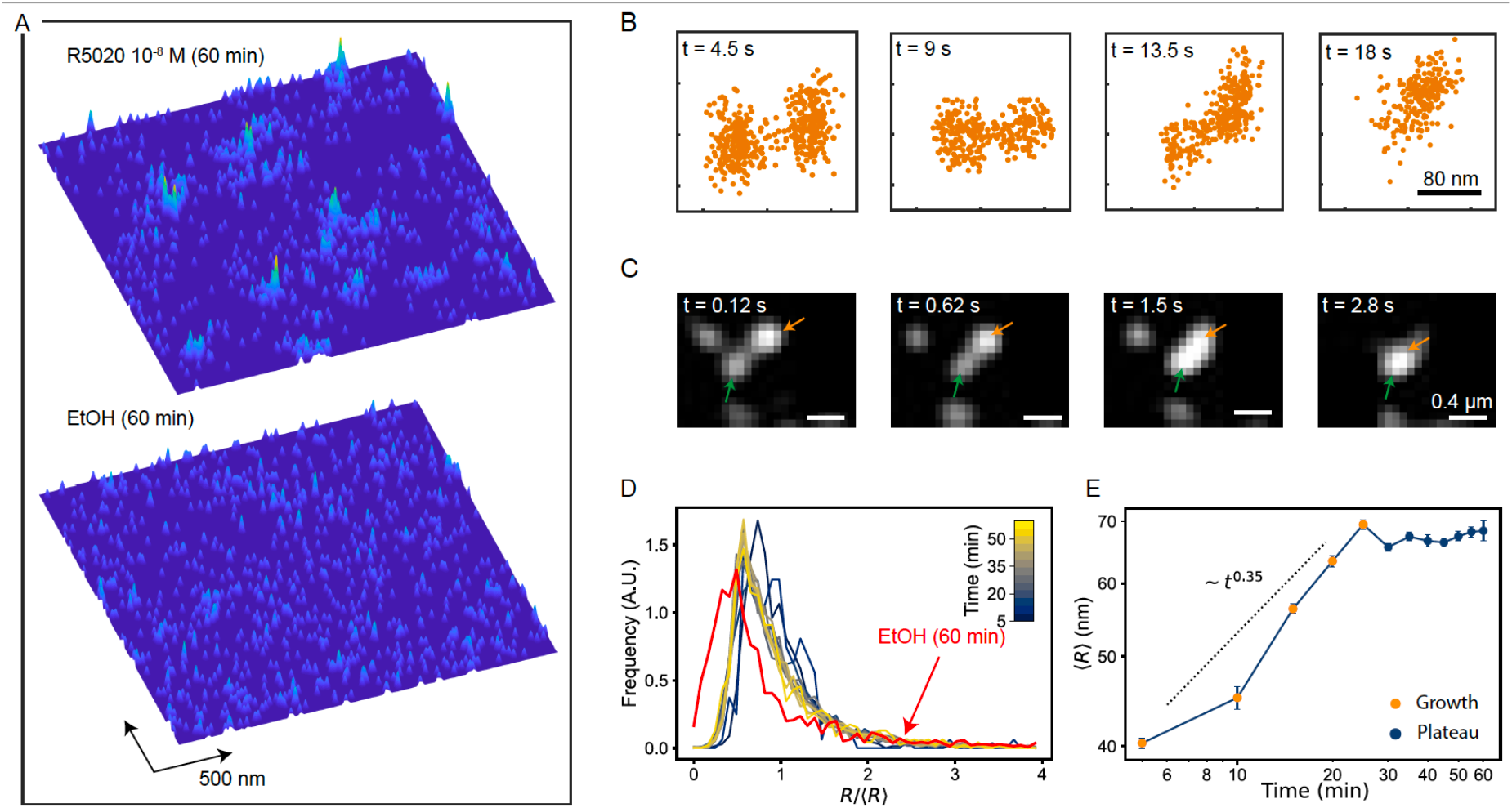
Nanometer-scale spatiotemporal mapping of PR in living nuclei. (A) 2D density maps of individual PR localizations collected over 75 s on an area of 2.4 × 2.4 μm^2^, after 1 hour of ligand stimulation (upper panel) and control (lower panel). Each map contains 1000 localizations. (B) Snapshots of two different condensates as they merge over the indicated time windows. The 2D maps have been generated by cumulating single molecule localizations in time windows of 4.5 s (300 frames). (C) Merging events of two different PR condensates (highlighted by orange and green arrows) visualized by confocal microscopy using GFP labeling conditions. (D) Distribution of PR condensate radius normalized to the mean radius, over a time course of 60 min after 10^−8^ M hormone stimulation. Each curve corresponds to a 5 min time point. The red curve corresponds to the size distribution in the absence of the hormone. (E) Mean condensate radius as function of time. At each time point, data correspond to several regions of interest (ROI) analyzed from two different cells and two separate experiments.

To enquire on the physical mechanism that leads to PR condensation in the nucleus, we first relied on the fact that the 2D density maps also contain temporal information. We thus accumulated localizations for time intervals of 4.5 s (300 frames) to build up the temporal evolution of condensates during an observation time of 18 s, and used a cluster algorithm(40) (see Methods) to detect condensates formed by the local accumulation of individual localizations. We readily observed merging events of individual condensates in time (Fig. 3B), which were also confirmed by confocal video imaging at high temporal resolution of fully saturated GFP-labeled PR molecules (Fig. 3C). The merging of condensates is a first indication that its growth is dominated by a BMC process.

Since PR condensation in our system can be accurately tuned by the time and amount of hormone addition, we took advantage of this property to further assess the condensate growth mechanism. For this, we generated 2D density maps of single molecule localizations over a time course of 60 min starting right after adding the hormone (10^−8^ M). We cumulated localizations over 5 min intervals and used the cluster algorithm to generate distributions of condensate radii at each 5 min time point. Interestingly, we found a log-normal distribution of the condensate radii (Fig. 3D), similar to that described for systems undergoing LLPS under a BMC mechanism(24). Note that a similar size distribution was also observed in the absence of hormone (EtOH), indicating a preexisting population of small condensates, in agreement with our SPT data. In addition, we calculated the mean radius size of the condensates over time (Fig. 3E). Two distinct regimes can be clearly identified. During the first 30 min, the average radius grows following a power law, <*R*> ∼ *t*^*β*^, with a fitted *β* = 0.3505. After 30 min, the system reaches a steady-state plateau in which the average size of condensates remains constant. The initial growth scaling exponent, the log-normal distribution of the condensates’ radii as well as the presence of merging events, are all consistent with a BMC-based condensate growth mechanism(41).

Intriguingly, whereas the classical BMC model predicts that condensates grow in time until forming a single droplet(41), our system clearly deviates at longer times from such prediction, reaching a plateau with condensates sizes around 70 nm (Fig. 3E). To understand such a nanoscale arrested growth, we took a closer look to our SPT data. Despite the short length of the trajectories, we could readily detect the occurrence of escaping events, i.e., particles being able to exit the condensate (Fig. S5). Such escaping behavior has been also recently observed on DNA repair condensates in living cells(42). These observations suggest that particle escaping could influence PR condensate growth at the steady-state in the nucleus.

### Particle escaping leads to nanoscale arrested growth of PR condensates

To investigate if the presence of escaping events in a BMC scenario could lead to an arrested growth of condensates with a plateau on their sizes, we developed a theoretical model in which particles —PR dimers in our case, or other biological components in a general context— diffuse freely through the system, but also interact with each other in a non-trivial way. The model is based on the main principles of BMC: when two particles coincide (i.e., they contact each other), they interact together forming a condensate. Subsequent new interactions make the condensates to grow until reaching a phase separated system in which all the particles segregate from the environment forming a single condensate. To include the effect of particle escaping events in our model, we simulated a system of particles performing BMC-like condensate growth but incorporating a probability, *P*_*u*_, that particles escape from condensates (see Methods). Compared to the classical BMC model (Fig. 4A, upper panels), the presence of escaping events (i.e., *P*_*u*_> 0) prevents the system from reaching the single condensate state (Fig. 4A, lower panels) resembling our experimental observations.

**Figure 4.**
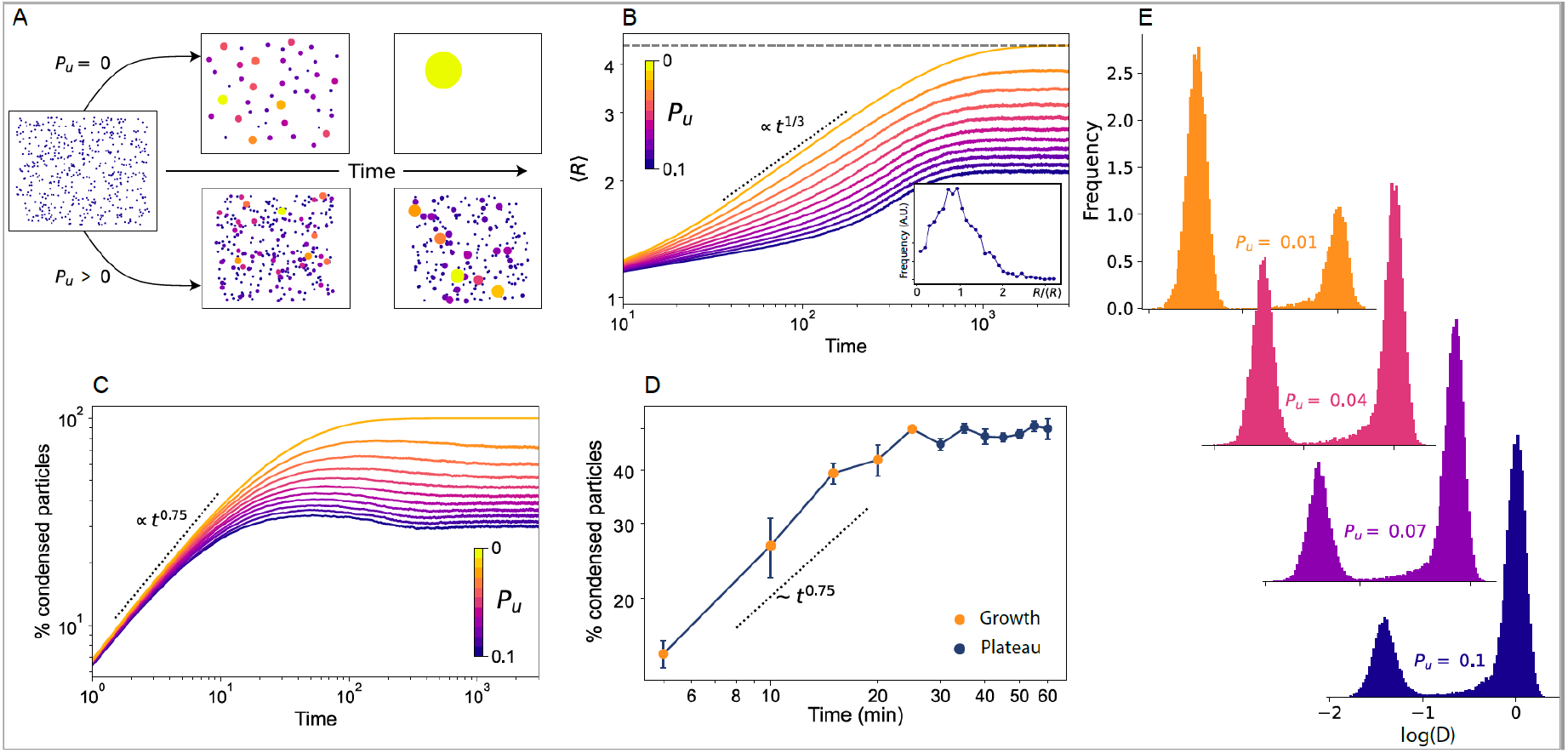
Extended BMC model including stochastic unbinding of PR molecules from condensates. (A) Snapshots of two simulations of the theoretical model, showcasing the temporal evolution of two systems, one with *P*_*u*_= 0 (top) and one with *P*_*u*_> 0 (bottom). (B) Mean radius size evolution as a function of time, for a system of *N* = 20, *L* = *N*/0.05. Each color represents the result for a different *P*_*u*_. The dotted line shows the expect BMC growth (<R> ≈ *t*^1/3^). The horizontal dashed line shows the maximum mean size possible for the simulated system (<R> =√*N*). The inset shows the steady-state normalized radius distribution for a system of *N* = 500 and *L* = *N*/0.01 for *P*_*u*_= 0.2. (C) Percentage of particles forming condensates as function of time for different *P*_*u*_values. (D) Experimental data showing the percentage of particles forming condensates as function of time. The data corresponds to the same experiments shown in Fig. 3D, E. (E) Diffusion coefficient distributions resulting from the simulations, for free particles (centered around Log(*D*) = 0) and of condensates (left distribution) for four different *P*_*u*_values.

We performed simulations considering that at each time step particles have a probability *P*_*u*_of unbinding and exiting the droplet in which they are contained, and calculated the average size of the condensates <*R*> as a function of time (Fig. 4B). For BMC (namely *P*_*u*_= 0), the system grows following the expected relation <*R*> ≈ *t*^1/3^ with a final single condensate size (dotted line). For values of *P*_*u*_> 0, condensate growth follows the same power law scaling, but notably, the system reaches a plateau with a steady-state mean radius <*R*> *∞* of smaller size, akin to our experimental observations. The larger the escaping probability, the smaller the final radius of the condensates (Fig. S6). Using the simulations we also generated distributions of the steady-state condensate sizes for different *P*_*u*_. An example of such distribution for *P*_*u*_= 0.2 is shown in the inset of Fig. 4B, exhibiting the expected log normal distribution for BMC processes. Importantly, the steady-state size distribution derived from the simulations is qualitatively similar to that obtained from our experimentally generated 2D density maps (Fig. 3D).

To further validate our model in terms of predictions that could be experimentally tested, we calculated from the simulations the percentage of condensed particles as function of time, for different *P*_*u*_. For a standard BMC process (*P*_*u*_= 0), in the state-state regime a single condensate will be formed, and accordingly, the percentage of condensed particles should reach ∼ 100%. However, in a scenario in which particles have a certain probability from escaping a condensate, the balance between coalescing and escaping events should maintain the percentage of condensed particles constant after the initial growth period. Our model predicts that the percentage of condensed particles increases as *t* ^0.75^ for shorter times and reaches a plateau with a constant percentage of condensed particles, whose value is again dependent on *P*_*u*_(Fig. 4C). To experimentally test this prediction, we extracted the percentage of condensed particles from our experimental 2D density maps at different ligand exposure times (10^−8^ M hormone concentration). Remarkably, our experimental data show an increase in particle condensation at early growth times with an exponent similar as to the one predicted by our model, and most importantly, it also exhibits a plateau in the percentage of particles forming condensates after 30 min of hormone exposure (Fig. 4D). Hence, our model of BMC-condensate growth together with condensate particle escaping is able to recapitulate our experimental data and to make predictions fully testable at the single molecule level in living cells.

Finally, we generated diffusion coefficient histograms from our simulations. As our experimental data are consistent with a BMC mechanism (i.e., condensate growth ∼ *t*^1/3^) we considered the presence of Stokes drag, as it is usual in a BMC process. Hence, free particles (i.e., outside condensates) would diffuse with diffusion coefficient *D*, while condensates of size *R* would diffuse with diffusion coefficient *D*_*r*_= *D*/*R*. Moreover, to account for the heterogeneities present in any biological scenario, we added a small random noise to each *D* value. We generated *in silico* distributions of the *D* values at steady-state, for various *P*_*u*_(i.e., accounting for the final condensate radius *R* and the percentage of free *vs*. condensed particles at each given *P*_*u*_). As expected, Fig. 4E shows the appearance of two distinct distributions, with a peak at *D* = 1 corresponding to the diffusion of free particles and a second peak at lower *D*, which is an effect of the Stokes drag and hence corresponds to the condensates diffusion. Interestingly, decreasing *P*_*u*_effectively increases the number and sizes of the condensates and reduces the number of free particles, resulting in an similar effect as to the increase of hormone concentration. Based on these results we suggest that at low hormone concentrations, the escaping probability of PR molecules from small condensates is large, leading to a large number of free, non-condensed particles. As hormone concentration increases beyond a critical value, the PR escaping probability reduces so that condensates grow reaching a finite stable size which is ultimately controlled by *P*_*u*_.

## Discussion

We have presented a single molecule study of the physical properties of transcriptional condensates *in vivo*. The inducibility of our system to undergo phase separation by means of hormone concentration and exposure time has allowed us to accurately tune the onset of phase separation as well as to thoroughly investigate the growth dynamics of nuclear PR condensates in living cells. Interestingly, we found that while growth dynamics of PR condensates is dominated by BMC at shorter times, condensates exhibited arrested growth reaching nanoscale sizes at longer timescales, clearly deviating from a classical BMC mechanism. To rationalize our results we proposed an extension of the BMC model by including stochastic unbinding of particles within condensates, i.e., introducing a probability that considers particle escaping from condensates. With this minimal consideration, our model fully reproduces the key features of an experimental system undergoing phase separation *in vivo*. Moreover, by modulating the probability of particle escaping, our model is able to predict the final condensate sizes, the population of molecules partitioning inside or outside condensates as well as their diffusion behavior. As a whole our experimental data and theoretical model are consistent with droplet growth dynamics being ultimately controlled by the escaping probability of transcription factors molecules within condensates. The interplay between condensation assembly and single molecule escaping thus maintain a preferred and maximum physical condensate size. Particle escaping from condensates can account for an exquisite control of the condensate size in non-equilibrium systems such as the cell, as also recently observed in other biological scenarios such as DNA repair condensates(42). This mechanism might provide a delicate fine-tuning by the cell that prevents a single phase that would lead to transcription collapse or chromatin condensation.

Recent SPT experiments showed that TFs transiently bind to DNA with rather short binding times (in the seconds scale)(13, 14). We propose that condensate formation might increase the likelihood that individual PRs rebind within short timescales to their corresponding DNA binding region. Such a condensate environment will thus increase the effective time that a given DNA region is bound by TFs. This hypothesis is further substantiated by our experimental data analyzed by machine learning, where FBM, traditionally associated to diffusion within viscoelastic media, was found to describe best PR low mobility diffusion. In conclusion, the exclusive combination of single molecule sensitive imaging techniques together with theory and simulations as reported here, brings a substantial step forward in understanding the behavior of individual proteins within condensates.

TF condensation has been customary studied through ensemble or static measurements, mostly in *in-vitro* settings or in fixed cells. In contrast, the experiments and theoretical model presented here provide a general framework to investigate the dynamics of phase-separation in living cells at the single molecule level. Moreover, our approach can be further extended to a wide range of biological systems as well as other soft-matter based interacting systems. Overall, this work brings unique insights into phase separation in soft matter systems from both experimental and theoretical perspectives.

## Materials and Methods

### Plasmids

The original pGFP-PRB was a gift from Gordon Hager (National Cancer Institute, NIH, Bethesda, USA). This plasmid expresses the PR isoform-B under a tetracycline controllable promoter (TetOff system, Clontech). To perform the SPT experiments, a SNAP tag was introduced at the N-terminal to the GFP, using Gibson cloning (pSNAP-GFP-PRB). A Puromycin resistance plasmid (pPUR, Clontech, Cat No. 631601) was used as a selection marker. All plasmids were linearized with ScaI before electroporation.

### Cell culture and electroporation

MCF7 Tet-off cells (Clontech, Cat No. 631154) were grown on Dulbecco Modified Eagle Medium (DMEM) high-glucose media supplemented with 10% Tet-free Fetal Bovine Serum, 2mM L-glutamine, 1 mM sodium pyruvate, 100 U mL-1 penicillin and 100 μg mL-1 streptomycin. The cells were cultured at 37°C in a CO_2_/air (5%/95%) incubator. Cells were electroporated simultaneously with the pSNAP-GFP-PRB and the pPUR, using a 10 to 1 ratio respectively. Electroporation was performed using the Amaxa Cell Line Nucleofector Kit V (Lonza) using the P-20 program, following manufacturer’s instructions. After one week, cells were selected under 0.6 μg/ml Puromycin, to enrich for electroporated cells, and then sorted in single cell wells using GFP as a marker, in order to generate a stable cell line.

### Hormone stimulation and SNAP labeling

Two days before the microscopy, around 200.000 cells were seeded in 35 mm glass bottom dishes. Sixteen hours before hormone stimulation, cells were washed with Phosphate-buffered saline solution, to eliminate traces of phenol red, and then changed to white DMEM media supplemented with 10% charcoal-treated FBS Serum, 2mM L-glutamine, 1 mM sodium pyruvate, 100 μg mL-1 penicillin and 100 μg mL-1 streptomycin; from now on abbreviated as “charcoalized white DMEM”. The Janelia Fluor® 549 dye coupled to the SNAP substrate was kindly provided by Luke Lavis (Janelia Farm, Ashburn, Virginia, USA). Cells were incubated with 10 nM for SPT and 100 nM for 2D spatiotemporal maps of the SNAP JF-549 dye in charcoalized white DMEM for 30 min at 37°C. Subsequently the cells were washed three times with PBS, and then placed back in the incubator in charcoalized white DMEM for one hour washout at 37°C. After the JF549 SNAP labeling, hormone stimulation was done using R5020 (Promegestone) solubilized in ethanol, or control conditions with this solvent. To study the response to different concentrations of hormone, a series of dilutions were made freshly before the microscopy acquisition. Time course experiments were performed at a hormone concentration of 10^−8^ M and SPT tracking data were recorded at intervals of 5 min during a total observation time of 60 min.

### Experimental setups

SPT and 2D spatiotemporal density maps imaging were performed in a Nikon N-STORM 4.0 microscope system for localization-based super-resolution microscopy, equipped with a TIRF 100x, 1.49 NA objective (Nikon, CFI SR HP Apochromat TIRF 100XC Oil). The sample was illuminated by a continuous 561 nm laser line with a power of 30 mW before the objective in HILO-configuration. The emission fluorescence of the JF549 dye was collected through the objective and projected into an EM-CCD Andor Ixon Ultra Camera at a framerate of 15 ms. The pixel size of the camera is 160 nm. During imaging, the temperature was kept at 37°C by an incubation chamber. GFP confocal line scanning microscopy was performed in a Leica TCS SP5 II CW-STED microscope using a 63x Oil, 1.4 NA objective (Leica HC PL APO 63x/1.40 Oil CS), using a multiline Argon laser at 488 nm for excitation. The emission fluorescence was detected with a Hybrid detector (Leica HyD) in photon counting mode, using a 500– 550 nm filtering. The sample was kept at 37°C with 5% CO_2_by an incubation chamber. For Fig. 3C, images of 256 × 256 pixels were acquired with pixel size of 80 nm and dwell time of 9 μs. Scanning was performed at 100 Hz, acquiring consecutive frames every 125 ms. For the Supplementary Video 1, images of 322 × 200 pixels were acquired with a pixel size of 160 nm. Each frame in the movie has a total integration time of 15 s, and corresponds to the sum intensity projection from 100 images taken consecutively every 150 ms, scanning at 700 Hz.

### Data analysis

To generate SPT trajectories, the nuclear region was segmented in the GFP channel intensity using Fiji. Individual tracks inside the nuclear region were analyzed using Trackmate(43). Particle detection was performed with a Difference of Gaussians, with an expected diameter of 0.6 μm and sub-pixel localization. Detected particles were first filtered based on the Signal to Noise Ratio of the input image and then based on quality score. The particles retained were then linked using a simple Linear Assignment Problem (LAP) tracker, with a 1 μm linking distance, 1 μm gap closing max distance and gap closing of two frames. Only tracks with more than 10 frames were considered for the analysis.

To generate 2D spatiotemporal maps, the total single molecule localizations of JF549 labeled PR molecules were detected by a custom Matlab Software over 5000 frames (75 s) and projected into one single frame. Condensates were detected by applying a Density-Based Spatial Clustering of Applications with Noise (DB-SCAN) (40)over the entire frame with a threshold of 48 nm of interparticle distance and condensates containing a minimum number of particles of 5. The radius was extracted by considering the area of the condensate a circle. The percentage of free particles was estimated by the number of particles not detected within a cluster divided by the total number of particles within a given area. The escaping events analysis was performed by taking PR trajectories and detecting within each trajectory a cluster by the clustering algorithm. Only trajectories where there is clear escaping event where considered. Time-evolution 2D density maps were generated by cumulating localization positions every 4.5 s, corresponding to 300 frames, for a total duration of 18 s.

Given a trajectory whose two dimensional position (*x, y*) is sampled at T discrete, regular time steps *t*_*i*_, its time averaged mean-square displacement (tMSD) was calculated using(44):

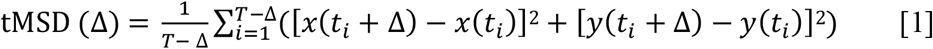

where Δ is usually referred as the time lag. Even in the presence of anomalous diffusion, at short times the MSD is well represented by

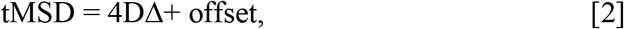

where *D* is the instantaneous diffusion coefficient. To extract *D*, we fit the tMSD between Δ = 2 to Δ = 4 and redefine it, as presented in main text, as *D*_2−4_.

For a given time, *t*, and a time between frames, δt, we define the turning angle, θ_*t*_, between consecutive trajectory segments, 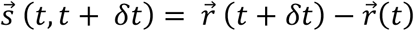, as follows (45):

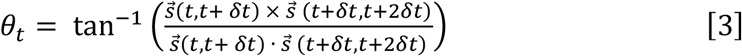

For our calculations, we consider the particle positions to be in 3D with the *z* component equal to zero. Using the above expression, the angles are defined between 0° and 360°. To calculate the anisotropy of the turning angles, the fold change between the number of angles from 180° ± 30° and 0° ± 30° was extracted(28).

### Machine learning architecture and analysis

A schematic pipeline of the machine learning (ML) method used in this study is presented in Fig. S2. The ML architecture is trained with a set of simulated trajectories, generate via the andi-datasets Python package(46). This tool allows to generate trajectories that are assigned to five different diffusion models. Moreover, trajectories with different anomalous exponents (0 < *α* < 1) can also be generated. The ML architecture can be trained separately to perform two different tasks: A) to classify the trajectories among a pool of different theoretical models; B) to regress the value of the anomalous exponent of each trajectory. Importantly, the training is done in a supervised way, i.e., we feed the trajectories to the machine, together with their corresponding labels (either the diffusion models for A, or the exponents for B). As architecture, we use a combination of gated recurrent units (GRU) and convolutional neuronal networks (CNN), merged with a contact layer made of fully connected neurons as depicted schematically in Fig. S2. The GRU layers are able to learn long-term features, while the CNN are a good strategy to tackle short length correlations(47). By combining the two approaches, we are able to characterize trajectories of only 10 data points in a robust manner.

In order to classify the experimental trajectories according to a given diffusion model, the last layer of the network consists in a soft-max layer of neurons, where is the number of models considered. The labels are encoded in a vector of elements, all equal to zero except the one encoding the model of the trajectory. The cost function to minimize is the Kullback-Leibler divergence which, for a set of trajectories 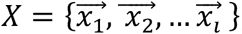, compares the output vector of the machine *f*_*m*_(*x*_*i*)_to the label vector 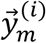 using

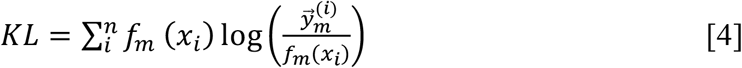

To faithfully characterize the set of experimental trajectories, we first train a model to classify among four diffusion models: continuous-time random walk (CTRW)(48), fractional Brownian motion (FBM)(37), annealed transient time motion (ATTM)(36) and the scaled Brownian motion (SBM)(49). For each model, we generated trajectories with anomalous exponent *α* ∈ [0.05, 1] in intervals of 0.05. We created a balanced dataset with 1000 trajectories per model and exponent, which in total sum up to 72000 trajectories. We separated the dataset into two, a training set with 57600 trajectories and a test set with 14400. The latter was used to calculate the accuracy of the model, i.e., to prevent the appearance of over-fitting. Note that the input size of the machine is fixed, which means that all the input trajectories should have the same size. As the experimental dataset has trajectories of varying size, from 10 to 1000 points, we solve such problem by restricting them to 20 frames long. This procedure ensures that most of the trajectories are considered, while the length is sufficiently large for the machine to have good accuracy. The trained model then has a micro-averaged F1-score of 0.733. When applied to the experimental dataset, 90% of the trajectories were classified either as FBM or ATTM.

Since the vast majority of the trajectories were classified either as FBM or ATTM, we trained the machine only with these two models. This allows to increase the accuracy of the ML classification for 20 frame long trajectories. In this case, the F1-score attained is of 0.822 (compared to 0.733). The confusion matrix for this classification is shown in Fig. S3A. The results of the prediction on the experimental dataset are presented in the main text.

For the anomalous exponent prediction, the output of the machine is a continuous value. Hence, the last layer of the neural network is a single neuron with a rectifier activation function (RELU). The loss function in this problem is the mean absolute error,

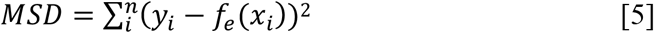

where *y*_*i*_is the label corresponding the trajectory *x*_*i*_, and *f*_*e*_(*x*_*i*)_is the network prediction. The sum is done over the set of trajectories in the training dataset. In order to infer the anomalous exponent for each individual trajectory, we used a simpler version of the neural network, containing 2 GRU layers of 100 and 50 neurons each, whose output enters two fully connected layers of 64 neurons and sigmoid activation functions. The last layer contains a single neuron with RELU activation function. Between each fully connected layer, we proceeded with a 25% dropout. This network shows a mean absolute error of 0.229 for trajectories of just 20 points. Note also that the predictions of the network are biased to increase the exponent, as shown in Fig. S3B.

### Theoretical model and simulations

Our theoretical model is based on the main principles of BMC, but with the addition of stochastic unbinding of particles from already formed condensates. In our system, particles diffuse performing Brownian motion through the system, but also interact with each other in a non-trivial way. When two particles coincide (i.e. they contact each other), they interact together forming a condensate. In a classical BMC process subsequent new interactions make the condensates grow until reaching a phase separated system in which all the particles segregate from the environment forming a single condensate. However in our case and motivated by our experimental observations we include an unbinding probability *P*_*u*_, such that at any given time, particles can exit the droplet in which they are contained.

The simulations consider the following free parameters:

***N*:** total number of particles;

***r:*** effective radius of the particles. We consider that all particles in the system have the same effective size and that they have a circular shape. If two particles of size *r* are closer than a distance 2*r*, they coalesce. When the two particles coalesce, the total area is conserved, such that the resulting droplet has area 2π*r*^2^. Then, a droplet containing *M* particles has a total radius of 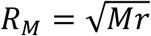 and area *A*_*M*_= *Mπr*.

***L:*** length of the 2D squared box acting as environment. The area of the box is hence L^2^. We consider in this case periodic boundary conditions, i.e., any particle traversing one of the borders of the box is immediately transferred to the opposite side. Similar simulations were performed with reflecting boundary conditions with analogous results.

***D:*** Diffusion coefficient of single particles. All particles (free as well as the condensate themselves) perform Brownian motion with the same diffusion coefficient *D*. Justified by the experimental results, as well as the theoretical considerations of BMC, we consider the presence of Stokes drag, i.e., a droplet of radius *R* will decrease its diffusion coefficient following *D*_*R*_= *D/R*.

***P***_***u***_: Unbinding probability. At each time step, particles have a probability *P*_*u*_of unbinding from the droplet in which they are contained.

***T:*** total number of time steps of the simulation.

The caption of Figure 4 contains the specific values of the parameters used for the simulations presented.

For simplicity, we usually consider *r* = 1 and *D* = 1. At the start of each simulation, all particles are distributed randomly, following a uniform distribution, all over the environment. The simulation then works as follows:

1. At the beginning of each time step, for every droplet containing more than one particle, we check how many particles unbind. Each particle has a probability *P*_*u*_of escaping from the droplet it is contain. All particles that have unbind will not be able to bind until the next time step (i.e., will not be considered in Step 3).
2. Each particle or droplet performs a spatial step, sampled from a Gaussian distribution of variance 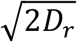; which effectively samples the steps of a Brownian particle with diffusion coefficient *D*_*r*_.
3. We iterate over each particle and droplet and find those who are in contact. These are consider to coalesce, forming larger droplets. We consider that the center of the resulting droplet is at the center of mass of the coalescing particles and droplets.
4. Repeat until doing T timesteps.

## Acknowledgments

We thank Gordon Hager for providing the pGFP-PRB plasmid and Luke Lavis for kindly providing the JF549 SNAP dye. We would like to thank the Advanced Light Microscopy Unit of the Center for Genomic Regulation (CRG, Barcelona) for their support. We thank G. Filion, R. Cortini, M. A. García-March and C. Manzo for fruitful discussions. The research leading to these results has received funding from BIST-Ignite funding (PHASE-CHROM) (to C.R. and J.A.T.-P.), the European Commission H2020 Program under grant agreement ERC Adv788546 (NANO-MEMEC) (to M.F.G.-P.), ERC AdG NOQIA (to M.L.) and ERC Synergy Grant 609989 (to M.B.), Government of Spain (Severo Ochoa CEX2019-000910-S, JdC-IJCI-2017-33160 (to J.A.T.-P.), State Research Agency (AEI) (FIDEUA PID2019-106901GB-I00/10.13039 / 501100011033 (to M.L.) and PID2020-113068RB-I00 / 10.13039/501100011033 (to M.F.G.-P.), QuantumCAT _U16-011424 (to M.L.), co-funded by ERDF Operational Program of Catalonia 2014-2020, QUANTERA MAQS (funded by State Research Agency (AEI) PCI2019-111828-2 /321 10.13039/501100011033), (to M.L.)); Obra Social La Caixa (LCF-ICFO) (to G.M.-G.), Fundació CELLEX (Barcelona), Fundació Mir-Puig and the Generalitat de Catalunya through the CERCA program and AGAUR (Grants No. 2017 SGR 1341 to M.L. and No. 2017SGR1000 to M.F.G.-P.).

## Supplementary Information

**Video S1. Time lapse of inducible PR nuclear condensates**. MCF7 cell-line expressing GFP-PRB before and after hormone stimulation. Before treatment with hormone the fluorescent signal of the GFP-PRB is homogeneous across the nucleoplasm. After hormone addition (R5020 10^−8^ M, black frames) the fluorescent signal distributes into condensates within 5 minutes of hormone exposure. Each frame has a total integration time of 15 s (see Methods).

**Figure S1.**
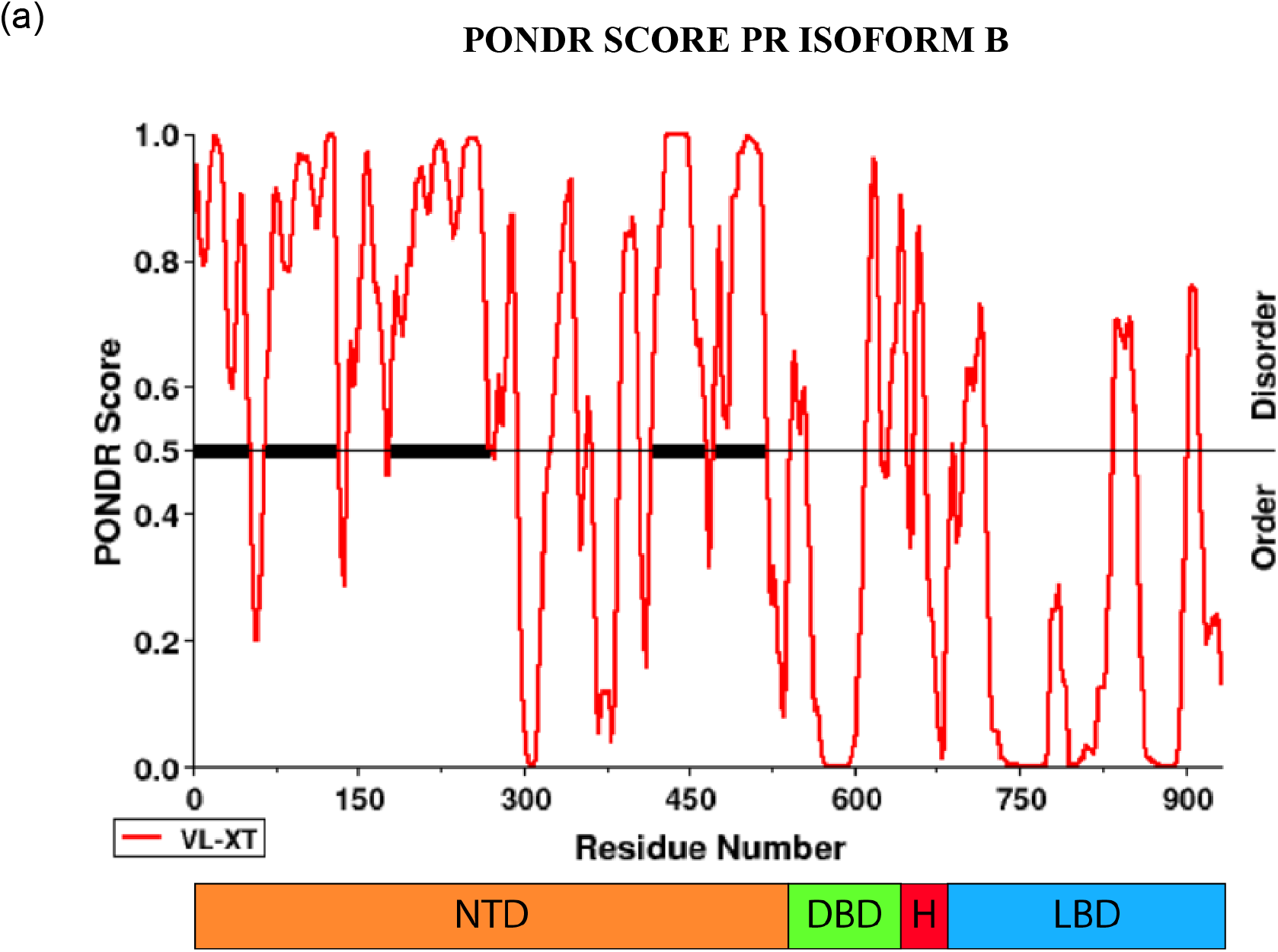
PONDR score of PR-B. (a) Prediction of Natural Disordered Regions (PONDR score) of PR-B generated at www.pondr.com. Note the different regions of PR-B denoted as N-terminal domain (NTD), DNA-binding domain (DBD), the Hinge (H) and the ligand binding domain (LBD). The NTD is highly disordered (PONDR score > 0.5)

**Figure S2.**
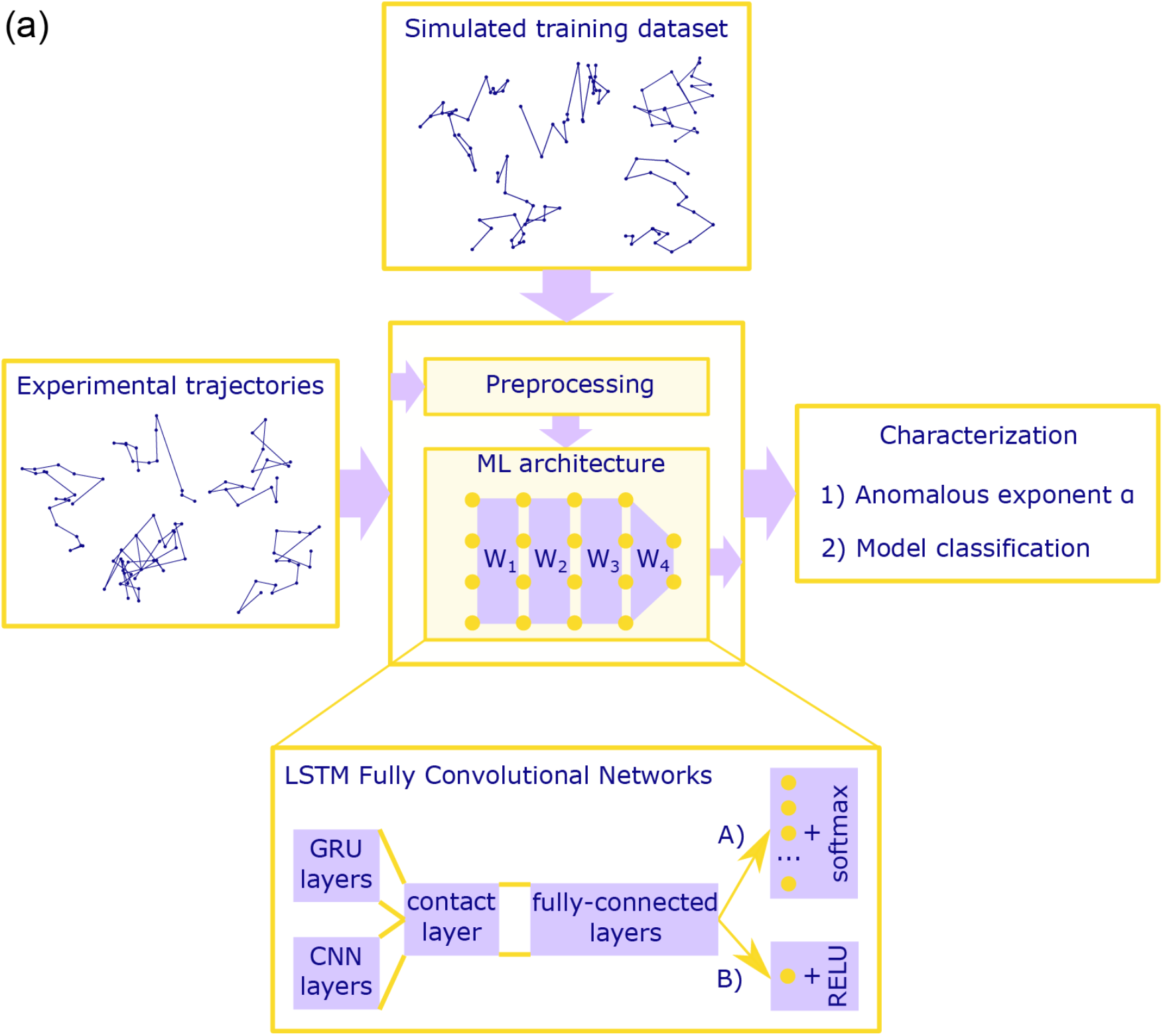
Scheme of the machine learning procedure. (a) The machine learning (ML) architecture is trained with a dataset consisting on simulated trajectories. Once the training is complete, the machine can assign to every experimental trajectory an anomalous exponent and a diffusion model.

**Figure S3.**
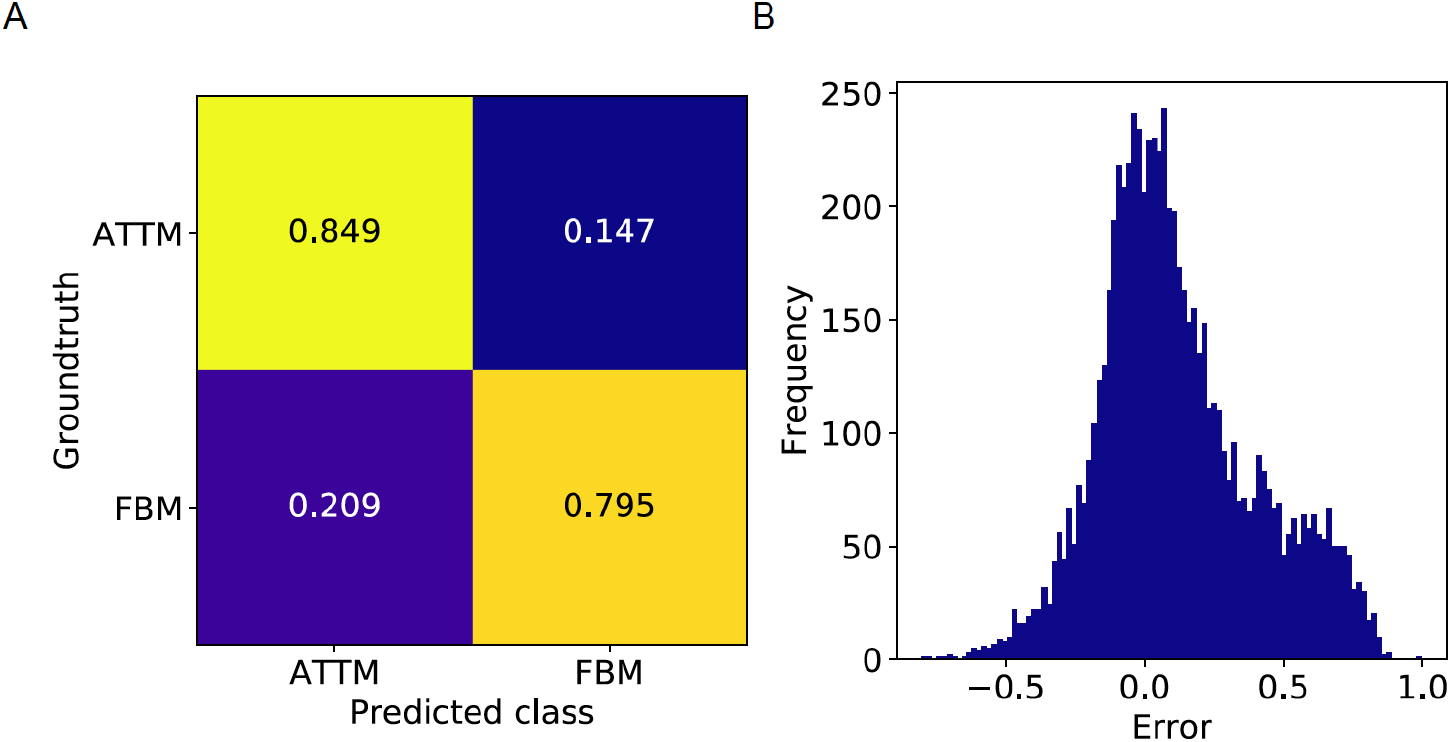
Error in the ML analysis. (A) Confusion matrix for the LSTM Fully convolutional network used for model classification. (B) Prediction error for the GRU network in the anomalous exponent prediction. For both cases, results were obtained using 7200 trajectories with T = 20, never seen by the machine, in order to avoid overfitting.

**Figure S4.**
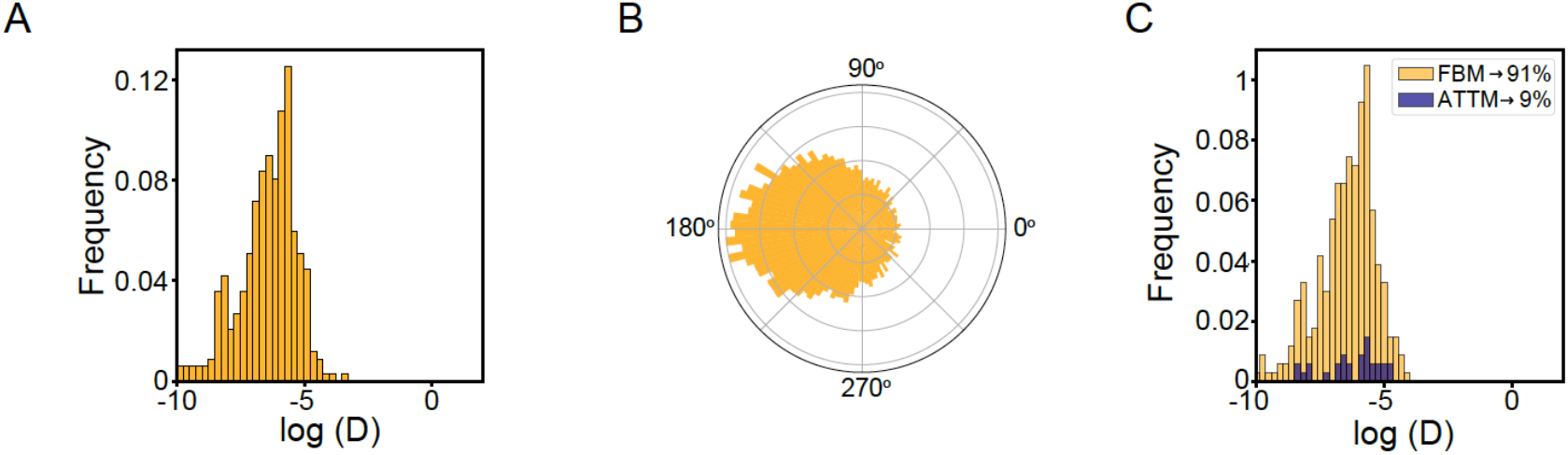
2D density maps confirms SPT data. (A) Distribution of the *D*_2-4_values of PR trajectories inside condensates and corresponding. (B) Angle distribution between successive steps, and (C) ML trajectory assignment to the diffusion behavior.

**Figure S5.**
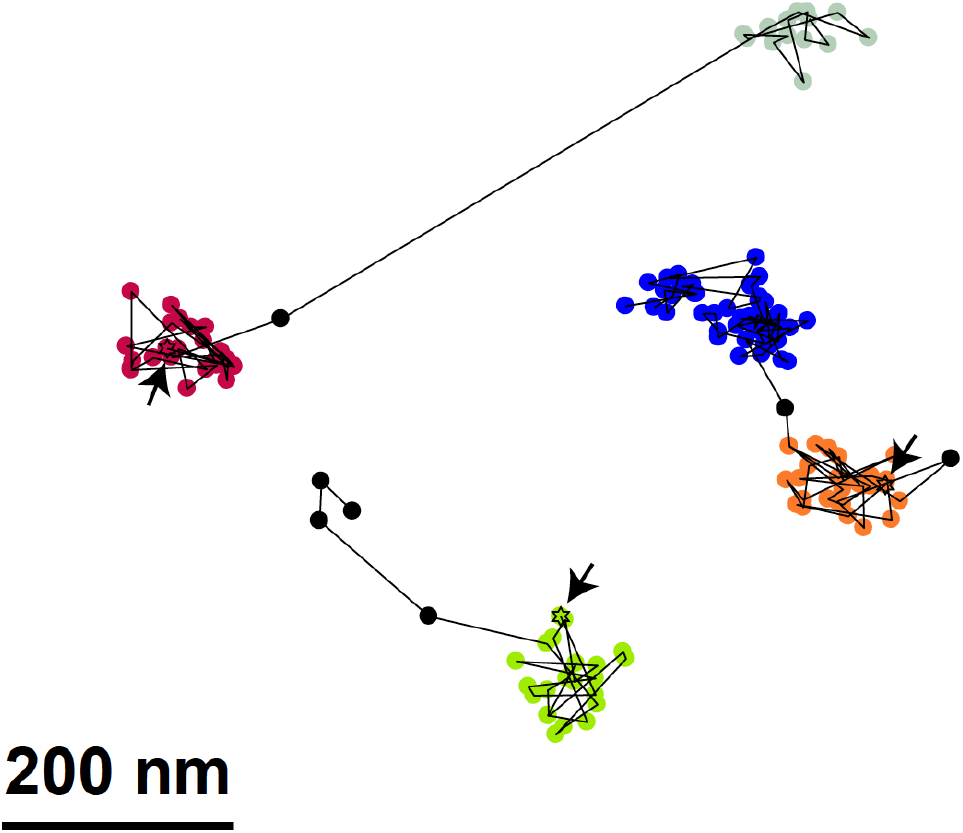
Escaping events in PR trajectories detected by the clustering algorithm. Representative trajectories with escaping events from condensates detected by the clustering algorithm. Condensates are labeled with different colors and the beginning of each trajectory is marked with an arrow.

**Figure S6.**
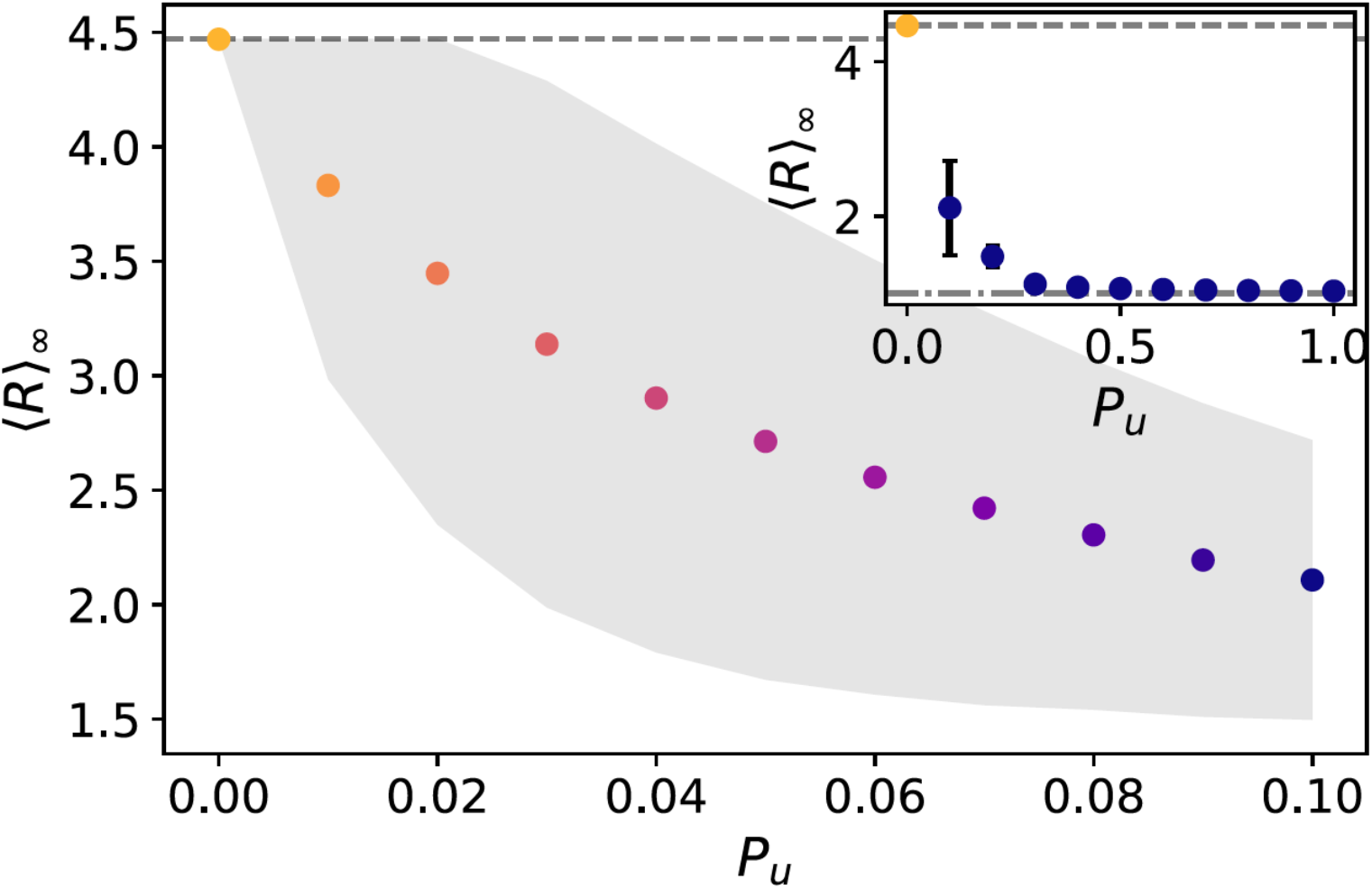
Steady state mean size radius as a function of the unbinding probability *P*_*u*_. Given is an average over 104 simulations. The gray region shows the variance between different simulations. The inset shows the complete range of *P*_*u*_, showing that above a certain *P*_*u*_, no condensates on average, are found.

## Notes

### Competing Interest Statement

The authors have declared no competing interest.

